# Multiomic Screening Unravels the Immunometabolic Signatures and Drug Targets of Age-Related Macular Degeneration

**DOI:** 10.1101/2024.05.07.592898

**Authors:** Xuehao Cui, Qiuchen Zhao, Bidesh Mahata, Dejia Wen, Patrick Yu-Wai-Man, Xiaorong Li

**Author notes:** Correspondence: Patrick Yu-Wai-Man, Xiaorong Li. Contributed equally (Xuehao Cui, Qiuchen Zhao).

## Abstract

Age-related macular degeneration (AMD) is a significant cause of visual impairment in the aging population, with the pathophysiology driven by a complex interplay of genetics, environmental influences and immunometabolic factors. These immunometabolic mechanisms, in particular, those distinguishing between the dry and wet forms of AMD, remain incompletely understood. Utilizing an integrated multiomic approach, incorporating Mendelian Randomization (MR) and single-cell RNA sequencing (scRNA-seq), we have effectively delineated distinct immunometabolic pathways implicated in the development of AMD. Our comprehensive analysis indicates that the androgen-IL10RA-CD16+ monocyte axis could protect against wet AMD. We have also identified several immune and metabolic signatures unique to each AMD subtype, with TNFα and Notch signaling pathways being central to disease progression. Furthermore, our analysis, leveraging expression Quantitative Trait Loci (eQTLs) from the Genotype-Tissue Expression (GTEx) project coupled with MR, have highlighted genes such as *MTOR*, *PLA2G7*, *MAPKAPK3*, *ANGPTL1*, and *ARNT* as prospective therapeutic targets. The therapeutic potential of these candidate genes was validated with observations from existing drug trial databases. Our robust genetic and transcriptomic approach has identified promising directions for novel AMD interventions, emphasizing the significance of an integrated multiomic approach in tackling this important cause of visual impairment.

## Introduction

Age-related macular degeneration (AMD) is a progressive retinal disease impacting the macula, which is the central segment crucial for precise vision (Fleckenstein et al., 2021). AMD ranks as a primary cause of vision impairment in individuals over the age of 55 across nations of various income levels, accounting for 6% to 9% of worldwide legal blindness (Mitchell et al., 2018; Wong et al., 2014). The global population affected by AMD is expected to increase from 196 million in 2020 to 288 million by 2040 (Wong et al., 2014). Early stages of AMD are identified through the presence of drusen, which are characteristic lipid-rich deposits found beneath the neurosensory retina (Murray et al., 2022). Vision impairment may develop at a later stage, either gradually due to geographic atrophy (GA, known as dry AMD or dAMD) or more swiftly through pathological neovascularization (known as wet AMD or wAMD). In cases of dry AMD, vision loss progresses slowly due to the atrophy of the retinal pigment epithelium (RPE), photoreceptors and choriocapillaris. Conversely, wet AMD is marked by the neovascularization from the choroidal vasculature of the eye, which is thought to be induced by increased expression of hypoxia-driven vascular endothelial growth factor A (VEGF-A), a process called choroidal neovascularization (CNV) (Guymer and Campbell, 2023). Wet AMD causes vision changes through CNV leakage, leading to fluid buildup, hemorrhages and fibrosis in and under the retina, defining it as exudative wAMD (Khachigian et al., 2023).

AMD is a multifaceted disease, intricately linked to aging, genetic predispositions, nutritional supplements, and environmental influences, stemming from a disruption in the retina’s homeostatic balance (Fleckenstein et al., 2024). A growing body of research highlights the immune system’s integral role in pathogenesis, with genome-wide association studies (GWAS) pinpointing polymorphisms in immune-centric genes, such as complement factor H (CFH), which elevate the risk of AMD (Klein et al., 2005; Strunz et al., 2020). This genetic susceptibility is further corroborated by pathological findings, showing an increased presence of immune cells, notably macrophages, in the choroid and/or Bruch’s membrane in AMD patient (Cherepanoff et al., 2010; Killingsworth et al., 1990; Oh et al., 1999). The recruitment of immune cells to CNV lesions, a key feature of the disease seen in animal models, indicates that the activation of macrophages and aging can affect disease progression positively or negatively (Manikandan et al., 2023). AMD’s complexity is amplified by the interaction of environmental and genetic factors, with more than 50 single-nucleotide polymorphisms (SNPs), predominantly related to the complement pathway and lipid metabolism, linked to an increased AMD risk (Fritsche et al., 2016; Han et al., 2020a). Furthermore, recent genetic studies have identified lipid and inflammatory biomarkers as indicators of AMD risk, underscoring the complex interplay between genetic makeup and environmental exposures in shaping disease trajectory (Fan et al., 2017; Han et al., 2020b; Han et al., 2021). Despite knowing that immune cells play a crucial role in AMD, the exact molecular signals guiding their activity are still unknown. This gap in knowledge has hindered the development of treatments for dry AMD, which makes up 90% of cases, limiting our ability to stop the disease from worsening. Therefore, understanding the immunometabolic pathways and genes involved in AMD is crucial to developing targeted new treatments and ultimately, improve patient outcomes.

Mendelian randomization (MR) can be used to make causal inferences in complex relationships. In MR, genetic variants associated with the exposure of interest are considered as instrumental variables to proxy for the exposure, and their association with an outcome is identified (Burgess et al., 2015). Single-cell RNA sequencing (scRNA-seq) have allowed for comprehensive analysis of the immune response and metabolic pathways (Papalexi and Satija, 2018). In this study, we have applied an integrated multi-omic approach to comparatively analyze immunometabolic signatures and related genes of AMD (summarized in Figure 1). We have leveraged data from GWASs of peripheral blood immune cells/traits (Orrù et al., 2020), inflammatory proteins (Zhao et al., 2023), metabolites (Chen et al., 2023), and various subtypes of AMD, early, dry, wet and mixed (Table S1).

**Figure 1.**
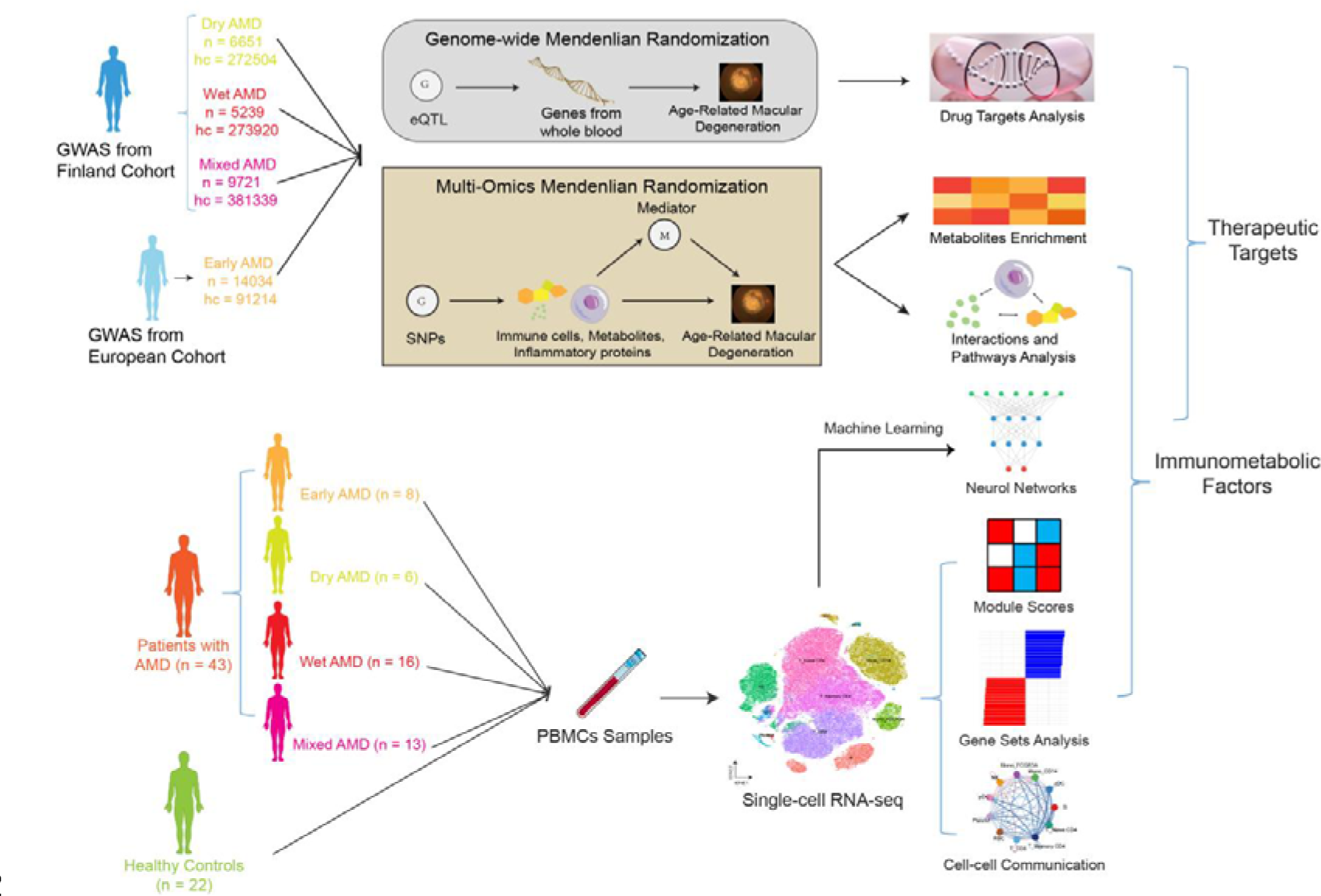
Study overview. An overview of this study’s data sources, analytical flow, and methodology. We performed multi-omic MR, scRNA-seq, genome-wide MR combined with metabolites enrichment analysis, mediation MR analysis, ML neurol network, cell-cell communications, gene sets analysis, drug targets analysis to explore and identify the immunometabolic pathways and drug therapeutic targets of AMD subtypes.

By capitalizing on integrated multiomics MR, we have uncovered how immune cells, inflammatory proteins, and metabolites are causally linked to AMD. By examining their interactions through mediation MR, we have identified critical immunometabolic pathways influencing AMD subtypes. Enrichment analysis of identified metabolites and scRNA-seq provided insights into the immune and metabolic changes in AMD, highlighting shifts in immune cell populations and metabolite profiles key to disease progression. We observed distinct cell-cell interaction patterns across AMD types, indicating unique immune responses per subtype. Machine learning (ML) on scRNA-seq data pinpointed genes differentially expressed in AMD, which were further investigated with MR analysis for their causal connections to disease pathophysiology. Finally, drug-target analysis explored new potential therapeutic avenues, aiming to pinpoint effective targets for future treatments (Finan et al., 2017).

Our study presents five major highlights: First, we found that several immune cells/traits and inflammatory proteins, particularly CD16+ monocytes and IL10RA, have a causal effect with AMD. Second, through metabolome-wide MR and scRNA-seq, we identified for the first time the importance of the androgen-IL10RA-CD16+ monocyte axis in wet AMD. Third, our analysis indicate that TNFα and Notching signaling pathways could play an important role in the progression of early AMD to CNV, and the bile acid (BA) metabolism pathway could play a significant role in dry AMD. Fourth, we found a series of genes associated with AMD subtypes and HLA-associated genes that demonstrated significant causal relationships in dry AMD and wet AMD. Fifth, our study also identified possible genetic drug targets based on significantly associated genes, including *MTOR*, *PLA2G7*, *MAPKAPK3*, *ANGPTL1*, and *ARNT*, and we have found a number of drugs that could potentially be used in the treatment of AMD.

## Results

### Study Design

An overview of the study design has been provided in Figure 1. First, we conducted multi-omics MR analyses to estimate the causal effects of immune cells/traits, metabolites and inflammatory proteins on AMD subtypes (Table S1). Second, we conducted enrichment analysis on metabolites with causal effects in MR and explored possible immunometabolic pathways using mediation MR and scRNA-seq. Third, the available scRNA-seq datasets were utilized to profile the fluctuation of peripheral immune populations as well as the transcriptional differences and cell-cell communication patterns between AMD subtypes and healthy controls. Fourth, we performed a genome-wide MR and ML to identify the genes associated with AMD. Ultimately, through the MR evidence and drug trial information, we identified some causal genes of AMD and potential drug targets.

In our study, inverse variance weighted (IVW), MR-Egger, weighted median, weighted mode and simple mode were used to estimate the causal relationship between genetically predicted exposures and AMD (Lee, 2020). IVW was used to identify 10 immune cells/traits, 10 inflammatory proteins and 19 metabolites with significant effects on AMD (Table1). In sensitivity analyses, we tested the robustness of the MR findings with a heterogeneity test, directional pleiotropy test and reverse-causation test to retain only the AMD subtypes that were robustly influenced by immune cells/traits, inflammatory proteins and metabolites (Table S14∼21). We did not find significant heterogeneity for the 10 immune cells/traits and inflammatory proteins with significant effects on AMD (*Q* > 0.05). The 10 immune cells/traits and 10 inflammatory proteins also showed no apparent sign of directional horizontal pleiotropy (Bowden et al., 2015) (*P* > 0.05). Many of these relationships were also confirmed across complementary MR methods except IVW (Table S22∼25). To assess potential reverse causation, we performed bidirectional MR that used AMD subtypes as the exposures and the immune cells/traits, inflammatory proteins as the outcome. None of these relationships were significant in the reverse direction (Table S18∼21).

### Multiomic MR Analysis Explores Causal Links Between Immune Cells/Traits, Inflammatory Proteins, and Metabolites in AMD

We treated 731 types of peripheral immune-related cells/traits as exposures and AMD subtypes as outcomes to explore whether the immune-related traits/cells would significantly affect AMD (Orrù et al., 2020). We identified 3 immune cells with significant effects on dry AMD, 2 cells positively impact dry AMD and a cell negatively impact dry AMD. We also identified 2 cells positively impact wet AMD with HLA DR on CD14-CD16+ monocyte is the most significant. Only unswitched memory %B cell was found that positively impact early AMD and in mixed AMD, we identified 4 immune cells all positively impact mixed AMD (Table 1). Our MR study indicates that a number of immune cells, particularly monocytes, along with their surface molecules and inflammatory activities, contribute to AMD. We plan to explore how immune cells influence AMD via specific pathways in future mediator analyses.

We performed a proteome-wide MR analysis of 91 blood inflammatory proteins on our AMD subtypes (Zhao et al., 2023). In dry AMD, the two most significant influences were observed from fibroblast growth factor 19 (FGF19) with a negative effect and signaling lymphocytic activation molecule (SLAM) with a positive effect. In wet AMD, VEGF-A and interleukin-15 receptor alpha (IL-15RA) both exerting positive effects, whereas interleukin-10 receptor alpha (IL-10RA) demonstrates a negative impact. There is no significant impact from inflammatory factors has been found in early AMD. In mixed AMD, 3 inflammatory proteins including VEGF-A can exert positive effects and FGF19, IL-10RA demonstrate negative impacts (Table 1). VEGF drives angiogenesis by binding with VEGFR2, initiating pathways for endothelial growth (Abhinand et al., 2016). Anti-VEGF treatments are the main approach for wet AMD, showing long-term effectiveness (Lanzetta et al., 2024). Our MR findings confirmed VEGF-A’s key role, validating MR is a trusted method to study AMD. Further, we examined the effects of inflammatory proteins on AMD through their role with immune cells and metabolites via mediation MR and scRNA-seq analysis.

### Metabolome-Wide MR Analysis Reveals AMD-related Immunometabolic Pathways

We conducted a metabolic-wide MR analysis that encompassed 1,091 blood metabolites and 309 metabolite ratios (Chen et al., 2023). In dry AMD, we identify 20 metabolites have a significant regulatory effect. In wet AMD, we found 4 metabolites; in mixed AMD, there were 6, while in early AMD, none were detected (Table1). We conducted heterogeneity and pleiotropy tests on the results and found that these results are free from pleiotropy (*P* > 0.05), although a portion of them demonstrated heterogeneity (*Q* > 0.05). For metabolites with heterogeneity, we employed the IVW method with a random-effects model to conduct MR analysis, thereby ensuring the accuracy of the results (Table S14∼21). Subsequently, we calculated the Z-scores (beta/se) for all significant metabolites at *P* -value < 0.05, positing that a positive Z-score indicates a stronger enhancing effect on AMD, while a negative Z-score suggests a greater inhibitory effect on AMD (Figure 2A∼D). In the AMD subtypes, we identified and marked the five metabolites with the most significant positive and negative regulatory effects.

**Figure 2.**
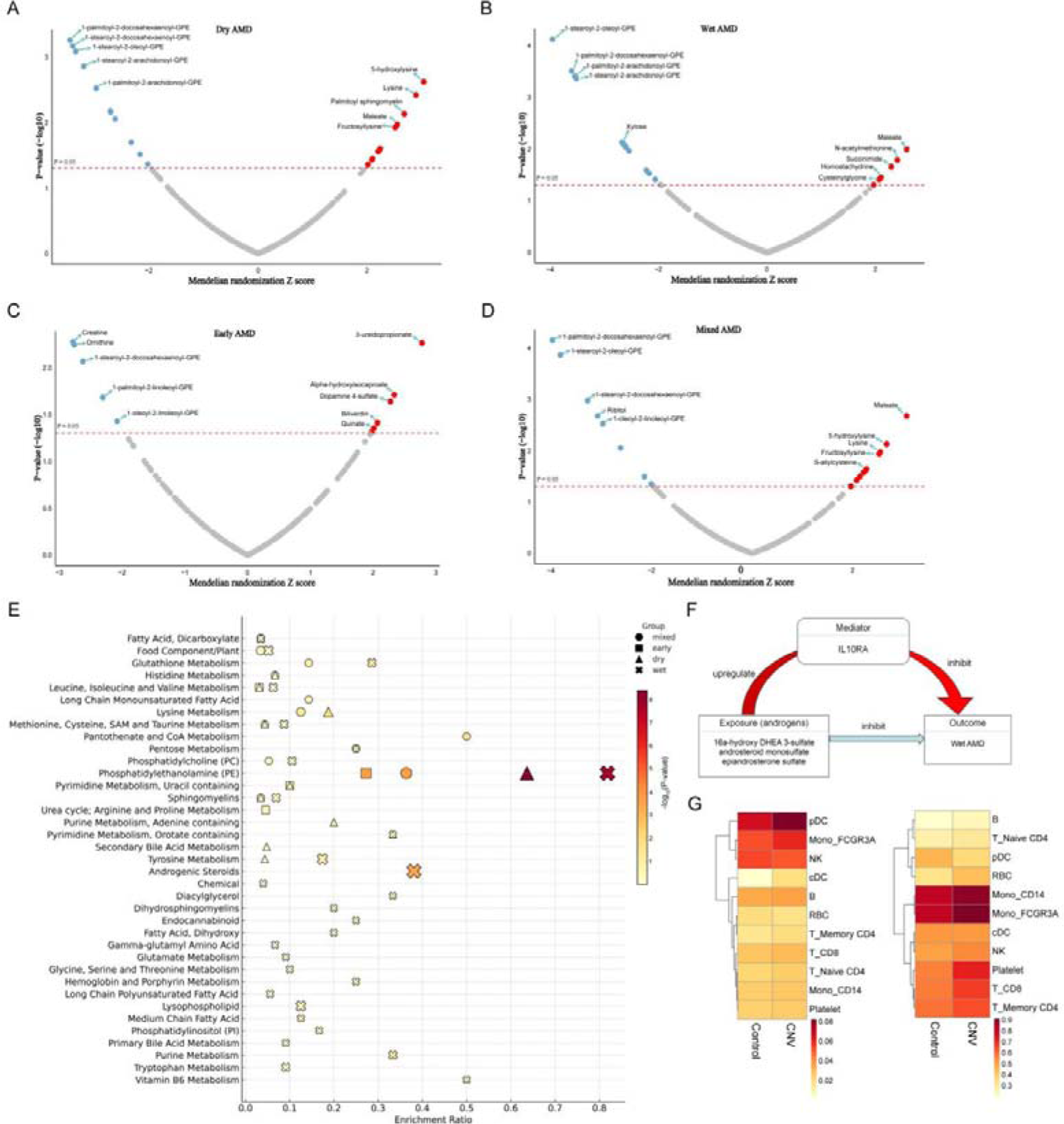
Metabolic MR, mediation MR and metabolic pathway analysis in AMD subtypes. A. The MR Z-scores for different metabolites in dry AMD with the horizontal axis representing the Z-score and the vertical axis indicating the p-value. Metabolites with positive effect are highlighted in red and negative are in blue. B∼D. The MR Z-scores for different metabolites in wet AMD (B), early AMD (C) and Mixed AMD (D). E. The enrichment analysis illustrates various metabolic pathways in AMD, with the size of the shape representing the number of hits in each pathway and the color indicating the group (Dry, Early, Wet, Mixed). The PE and androgen pathway among others show significant enrichment. F. This graph depicts the mediation effect that androgens upregulate the IL10RA, leading to the inhibition of wet AMD. G. The heatmap of the expression levels of various cell populations in control and AMD cases, with darker shades representing higher expression levels. The cell populations include monocytes, T cells, B cells, among others. The heatmap suggests differential cell type-specific involvement in AMD pathophysiology.

We performed enrichment analysis on metabolites with significant effects in the four AMD subtypes to explore which metabolic pathways play a role in the regulation of AMD (Figure 2E).

We discovered that the phosphatidylethanolamine (PE) metabolism pathway exhibits a highly expressed state across all four AMD subtypes, with particularly notable expression in wet AMD. Additionally, we observed that compared to the control group and other AMD subtypes, the androgen pathway exhibited significant expression in wet AMD.

We investigated the mediation effect of inflammatory proteins between wet AMD and androgen pathway-related metabolites: 16a-hydroxy DHEA 3-sulfate, andro steroid monosulfate, epiandrosterone sulfate, androstenediol (3β17β) monosulfate, and androstenediol (3β17β) disulfate (Figure 2F). We discovered that three metabolites associated with the androgen pathway: (1) 16a-hydroxy DHEA 3-sulfate, (2) androsteroid monosulfate, and (3) epiandrosterone sulfate—can suppress the development of wet AMD by upregulating IL-10RA. Additionally, epiandrosterone sulfate had the capacity to reduce IL-15RA, further contributing to the inhibition of wet AMD (Table S26). The mediation analysis revealed critical links between the androgen pathway, IL-10RA and wet AMD, highlighting the potential mechanisms and therapeutic target to modulate immune responses and curb disease progression. We conducted two sample MR analyses using different GWAS datasets of the androgen-IL10RA-CD16+ monocyte axis in the regulation of AMD, revealing a significant negative causal relationship between four types of androgens and wet AMD. Among these, one can upregulate the expression of the IL10 receptor. Concurrently, we found a significant negative causal relationship between IL-10 and wet AMD, thereby we double confirm the critical regulatory role of this immunometabolic axis in wet AMD.

Building upon the observed significant enrichment in the androgen pathway and its causal association with cytokine receptors such as IL10RA in wet AMD, we aimed to elucidate the differential responses of various immune populations to androgen and IL-10 signaling in both control subjects and CNV patients. Leveraging existing scRNA-seq datasets, we discerned a pronounced androgen response score predominantly in plasmacytoid dendritic cells (pDCs), CD16+ monocytes, and natural killer (NK) cells (Figure 2G). Notably, pDCs and CD16+ monocytes exhibited a heightened response in CNV patients. Regarding IL10RA expression, CD16+ monocytes demonstrated elevated levels compared to other cell types when juxtaposed with controls (Figure 2G). This finding corroborated our preceding MR analysis, suggesting that certain immune populations are intricately linked with metabolites and inflammatory mediators in the pathogenesis of wet AMD.

To further validate the causal relationship between the androgen pathway and IL10 in wet AMD, we selected several hormones directly related to the androgen pathway, along with IL10, as exposure factors, with wet AMD as the outcome factor for MR analysis (Table S37 ∼ S38). The results revealed significant negative causal relationships for 4-androsten-3β,17β-diol disulfate, androsterone sulfate, dehydroisoandrosterone sulfate (DHEA-S), epiandrosterone, and IL10 with wet AMD, indicating that these exposures play a protective role in wet AMD. Additionally, a significant causal relationship between 4-androsten-3β,17β-diol disulfate and IL10 suggests that it may further exert a protective effect in wet AMD by promoting the function of IL10.

### Systematic Immunoprofiling Reveals Cellular and Transcriptional Shifts in AMD Subtypes

Our prior analysis unveiled distinctive immunometabolic factors across various stages and types of AMD, prompting a systematic investigation into immune-related changes. We delved into the peripheral immune milieu at single-cell resolution in 43 AMD patients, encompassing 8 with early AMD, 6 with GA, 16 with CNV, 13 with mixed GA and CNV, alongside 22 healthy controls (Lin et al., 2024). Predominant cell populations were delineated utilizing scRNA-seq data (Figure 3A, S5A). Subsequently, cell fluctuations were quantified, illustrating the percentage deviation from controls across AMD subtypes (Figure 3B). Intriguingly, minor changes were observed between early AMD and controls, whereas more pronounced cell type fluctuations characterized advanced stages, notably an increase in CD8+ T cells in both GA and CNV, and a decrease in CD16+ monocytes in CNV (Figure 3B). Beyond cellular abundance shifts, we comprehensively mapped transcriptomic landscapes across 50 hallmark pathways for all groups under study (Figure 3C). This extensive profiling revealed a significant upregulation of the angiogenesis pathway in both mixed and CNV groups, aligning with their clinical manifestations (Heloterä and Kaarniranta, 2022). Furthermore, we identified unique signature pathways associated with specific AMD subtypes compared to controls, such as complement and interferon pathways in GA patients (Armento et al., 2021a). Additionally, the inflammatory response and TNFα signaling pathway were elevated in early AMD and CNV, highlighting potential prognostic targets for early AMD (Figure 3C). An inflammatory score was determined across different cell populations within AMD patients, pinpointing CD14+ and CD16+ monocytes as principal contributors (Figure 3D). Collectively, these findings underscore the presence of substantial cellular and transcriptional alterations across AMD subtypes, underscoring their potential as biomarkers and therapeutic targets.

**Figure 3.**
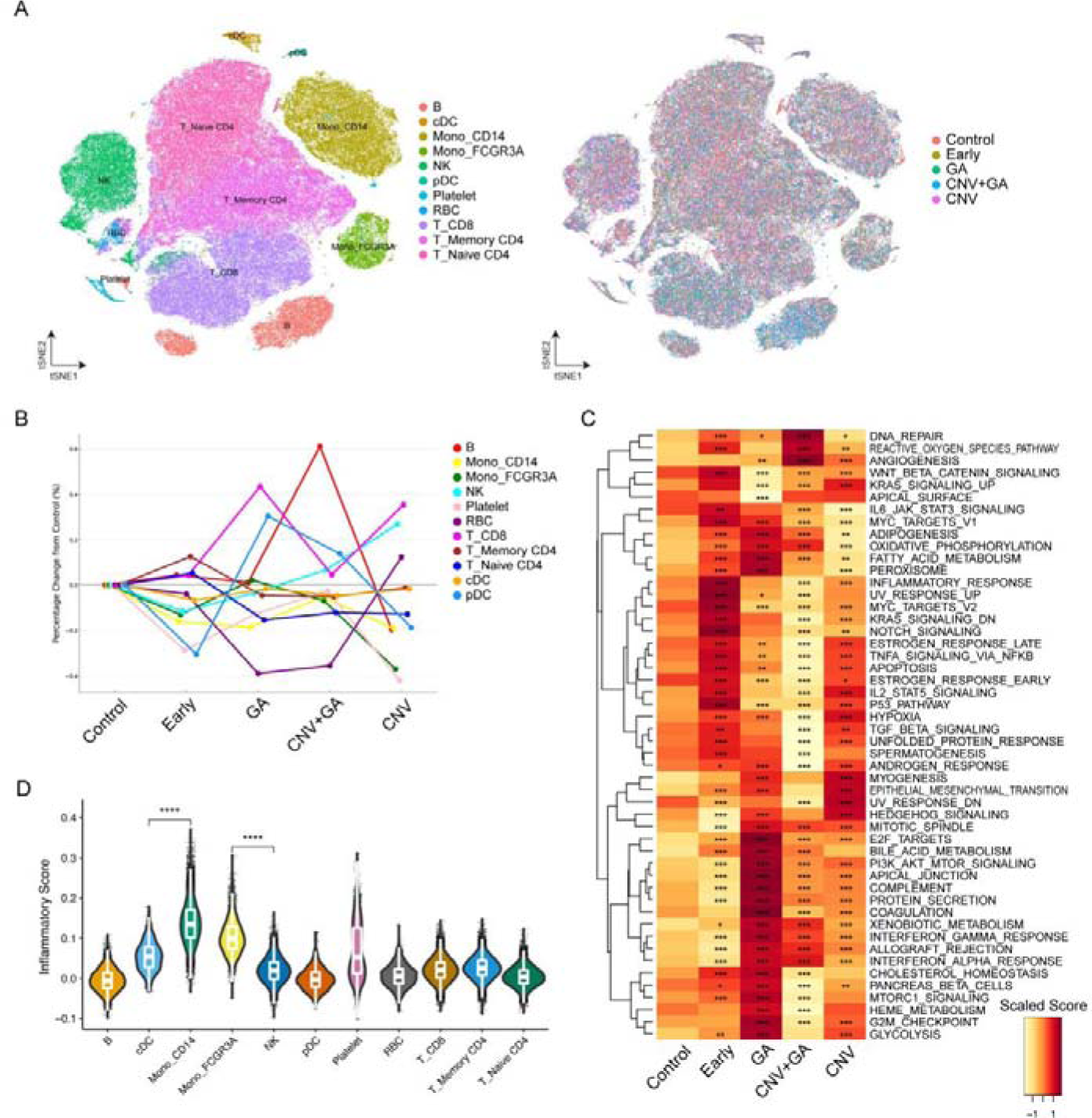
Systematic Immunoprofiling Reveals Cellular and Transcriptional Shifts in AMD Subtypes. A. t-SNE plots representing the distribution of immune cell populations from 43 AMD patients and 22 healthy controls. Different colors on the left highlight distinct cell subsets, while the right contrasts cell origins from different AMD subtypes and healthy controls. B. Bar graph detailing the percentage deviation of immune cell types across early AMD, geographic atrophy (GA), choroidal neovascularization (CNV), mixed GA and CNV, compared to healthy controls. C. Heatmap showcasing the gene set signature scores across 50 hallmark pathways for AMD subtypes and healthy controls. D. Violin plots illustrating the inflammatory score across various cell populations within AMD patients.

### Peripheral Immune Distinctions Between GA and CNV

In the light of the marked transcriptional difference observed across different types of AMD, our research extended into a detailed exploration of the peripheral immune landscape in patients with GA and CNV, two critical AMD subtypes. Utilizing Gene Set Enrichment Analysis (GSEA) on immune cells from these groups, we identified a pronounced reduction of CD4+ T cells, DCs, and macrophages activation signatures in CNV patients compared to those with GA (Figure 4A). Our subsequent analysis delved into the cell-cell communication dynamics, revealing that while the overall interaction landscape remained relatively stable between early AMD and controls (Figure S5B), there was a marked decrease in both the quantity and intensity of interactions within the CNV group compared with GA, especially among monocytes, dendritic cells, and T cells (Figure 4B, S5C). The assessment of relative signal flow further disclosed that pathways such as CCL/CXCL chemokines and interferon exhibited more robust signaling in GA, aligning with our earlier calculations of hallmark gene sets (Figure 4C). With a finer resolution, we highlighted changes in ligand-receptor pairs among the myeloid population and T cells that were significantly altered between GA and CNV (Figure 4D). This analysis showed a general decline in interactions within the CNV subgroup, especially between MHC-I/II and CD4/CD8 on cDCs/monocytes and T cells. Conversely, some interactions, such as TNF-TNFRSF1B between CD16+ monocytes and CD8+ T cells, were intensified in the CNV cohort, suggesting an elevated pro-inflammatory state (Figure 4D). Collectively, these findings underscore a disrupted peripheral immune response in CNV compared to GA, highlighting the nuanced immunological disparities underpinning these AMD subtypes.

**Figure 4.**
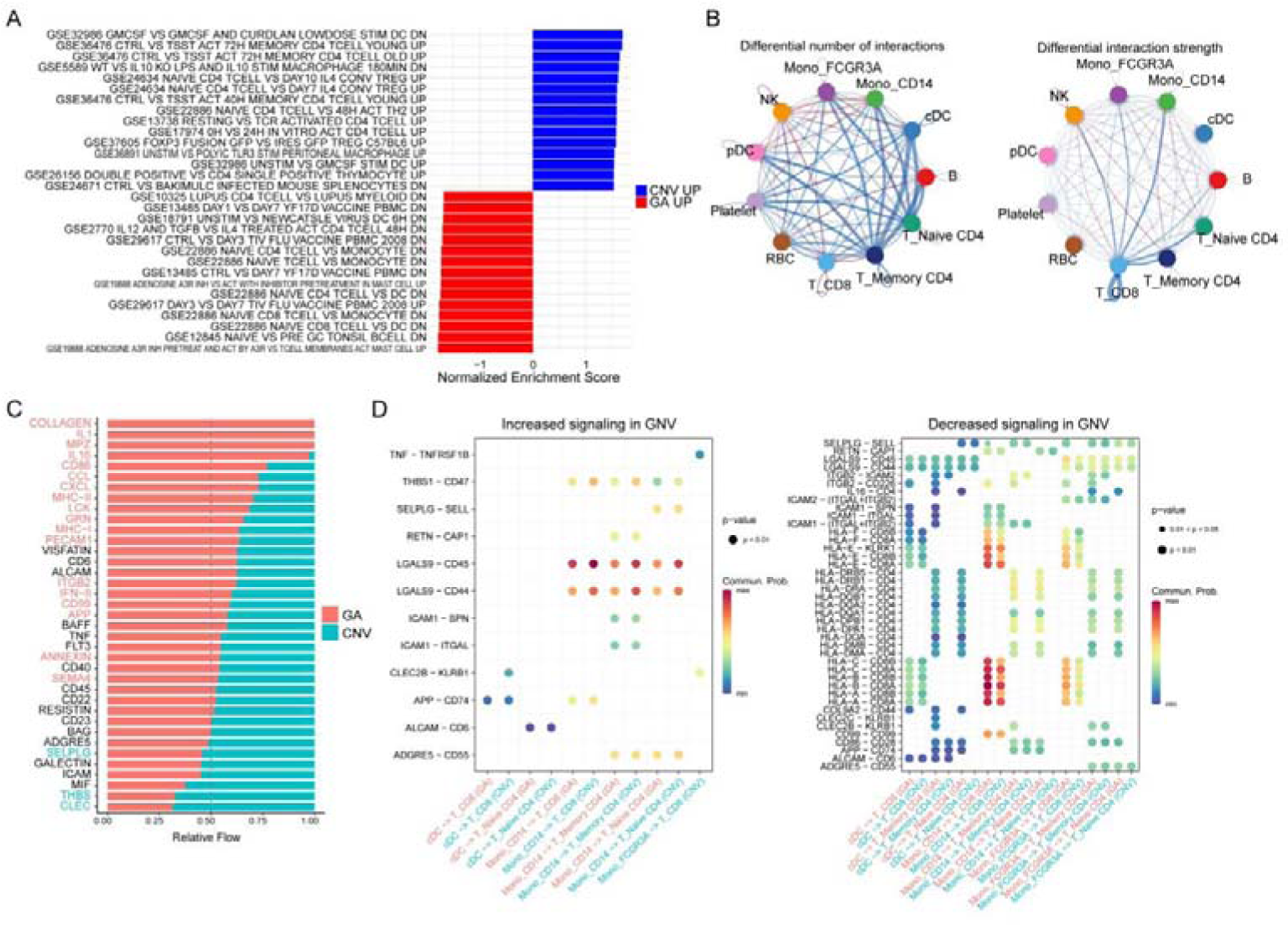
Peripheral Immune Distinctions Between GA and CNV. A. GSEA plots showing top 15 ImmuneSigDB gene set signatures between cells from patients with CNV and GA. B. Network diagrams showing cell-cell communication dynamics between peripheral cells in CNV and GA patient groups. C. Signal flow charts detailing pathway signaling differences between GA and CNV patients. D. Scatter plots of ligand-receptor pair expression in the myeloid population and T cells, comparing GA with CNV patients.

### Decoding Genes Linked to AMD Pathogenesis via Genetic Loci and Mediation MR Analysis

Upon identifying transcriptional variations between different AMD subtypes and healthy controls at single-cell resolution, we aimed to identify the feature genes related to AMD pathogenesis. Our study utilized expression quantitative trait loci (eQTL) from the Genotype-Tissue Expression (GTEx) project, which detailed genetic associations and gene expression across 49 tissues in 838 individuals, to identify genetic loci linked to AMD subtypes (Consortium, 2020).

We conducted MR analyses to assess the potential causal effects of gene expression levels on dry and wet AMD respectively. After MR analysis, we identified 31 potential causal genes with robust signals (*P* < 0.000015) in dry AMD and 29 potential causal genes in wet AMD (Table S32∼33). The gene enrichment analysis revealed that the genes associated with wet AMD were predominantly enriched in pathways related to leukocyte activation and immune response (Figure 5A), whereas those linked to dry AMD were primarily involved in the MHC class II protein complex assembly pathway (Figure 5B). Furthermore, we employed single-cell variational inference (scVI), which utilizes stochastic optimization and deep neural networks, on the scRNA-seq datasets from AMD patients and controls. This approach enabled us to precisely characterize the signature genes for both dry and wet AMD (Table S34∼36, Figure S6A, S6B). Interestingly, the causal association of *CXCL9, CX3CR1*, and *ZNF683* genes with wet AMD were doubly confirmed by both ML and MR methods (Figure 5C), suggesting a more significant causal link with wet AMD for these genes. Utilizing scRNA-seq data, we further mapped the expression levels of these three genes across all cell types from controls and AMD subtypes, and *CXCL9* was found to be highly expressed in CD16+ monocytes from wet AMD patients (Figure 5D). We also identified a significant association between 9 genes (*ACSL6, DDAH2, HLA-DQB1, HLA-DQB1-AS1, HLA-DQB2, HLA-DRB6, MEIKIN, MTOR,* and *PCCB*) and both dry and wet AMD, with four of these genes belonging to the Human Leukocyte Antigen (HLA) system, highlighting its potential role in AMD pathogenesis (Figure S6C, S6D). Further analysis using mediation MR based on multi-omic MR results revealed significant mediated effects of HLA-associated immune cells on AMD (Figure 5E). Specifically, the mediation analysis highlighted that 1-stearoyl-2-docosahexaenoyl-GPE, and lysine influenced the impact of HLA DR on immune cells in dry AMD, contributing to 4.58% and 10.9% of the effect, respectively, showing a subtle but important effect on immune response. For wet AMD, 1-palmitoyl-2-docosahexaenoyl-GPE helped slow the disease by affecting three types of immune cells through HLA DR, with effects between 6.42% and 7.85%, highlighting the complex relationship among specific lipids, immune cells, and HLA DR in AMD’s development (Figure 5E). Additionally, we noted a previously observed significant dysregulated communication between HLA-DR-CD4 in myeloid DCs or monocytes and CD4+ T cells (Figure 4D). These findings suggest a causal relationship between HLA-associated genes and immune cells in the development of AMD. In addition to HLA-related effects, our findings include CD4+ T cells mediating effects of TGF-A and CCL11 in dry AMD, and CD38 on CD20-B cells influencing FGF19’s effect (Figure 5E). For wet AMD, CD28+CD45RA+CD8br Treg cells counteract the disease via CX3CL1. A potential early AMD regulatory pathway involving CD127 on DN Treg cells suggests new prevention targets (Figure 5E). All in all, this comprehensive analysis elucidates the genetic underpinnings of AMD subtypes, suggesting novel targets for prevention and therapy.

**Figure 5.**
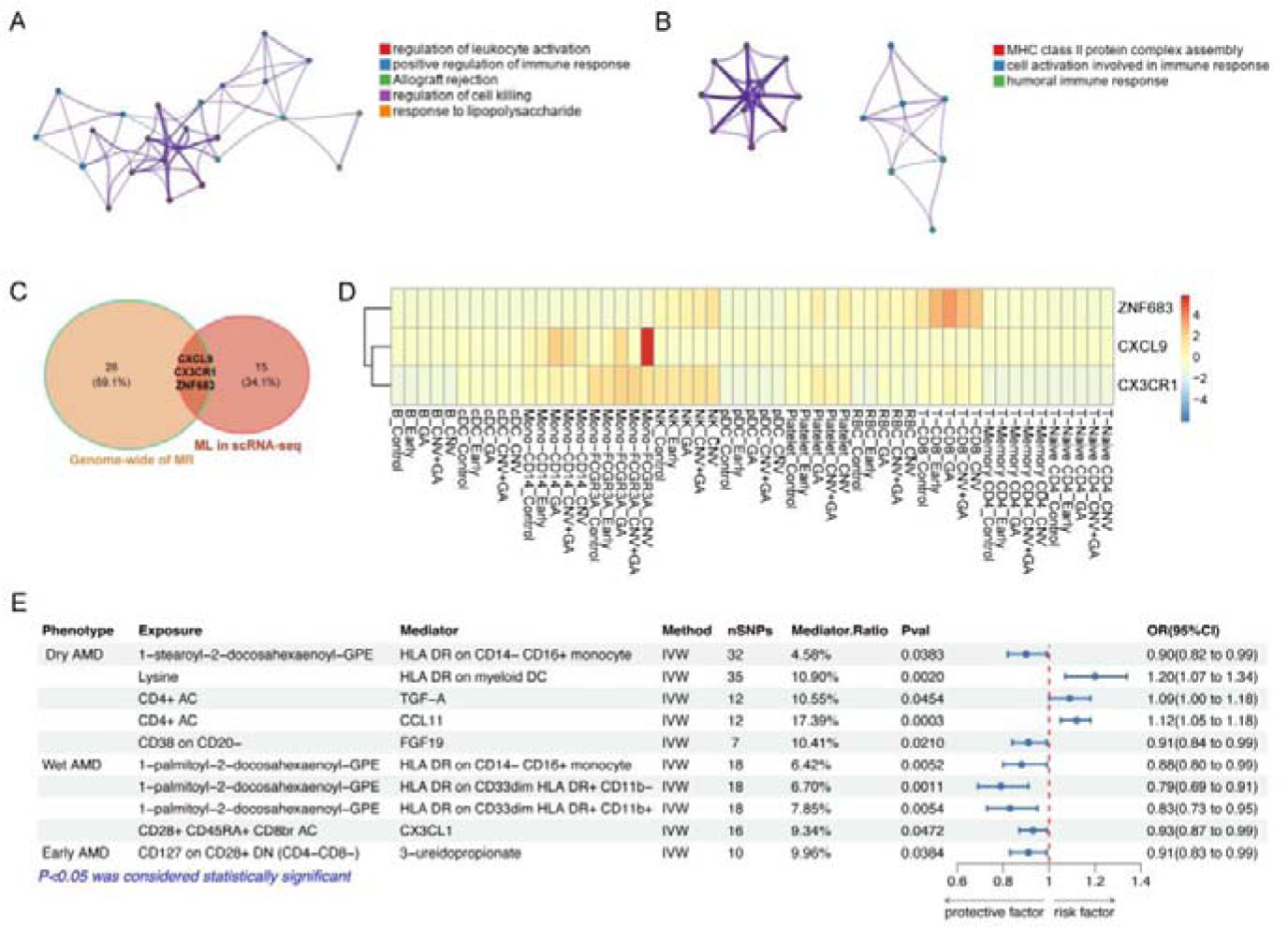
Integrated analysis of immune-related pathways and gene expression in AMD. A. The network illustrates the interactions between genes related to dry AMD, highlighting the key regulatory pathways such as negative regulation of immune response and regulation of leukocyte activation. B. The network shows the interactions of genes in wet AMD, with emphasis on MHC class II protein complex assembly and humoral immune response. C. The Venn diagram displays three genes associated with wet AMD that have been validated through both MR and ML methods. D. The graph shows the expression levels of genes in AMD and cells E. The graph summarizes the results of mediation MR analysis that identifies potential immunometabolic pathways related to AMD.

### Drug targets identification of AMD subtypes

Through our MR analyses, we identified 31 genes causally associated with dry AMD and 29 with wet AMD. To evaluate the potential of these genes as drug targets, we integrated MR findings with drug trial data from Open Targets and the Therapeutic Target Database, assessing the viability of these causal genes for therapeutic intervention (Table 2). First, genetically predicted higher expression levels of *MTOR* showed an effect on increasing the risk of both dry AMD and wet AMD. Some clinical trials supported the *MTOR* as a potential therapeutic target for wet AMD. Palomid-529 is a small molecular drug and a dual inhibitor targeting both *MTORC1* and *MTORC2* (Phase, 2012). A study reported that palomid-529 combined with pazopanib and bevacizumab were more effective in inhibiting the cell proliferation rates than any one anti-VEGF molecule alone in CNV (Mynampati Arunadithya and grover, 2021). In addition to *MTOR*, we have also identified *PLA2G7* as a potential drug therapy target, and drugs acting on this target, such as SNP318, may play a significant role in the treatment of dry AMD. Compared to dry AMD, besides *MTOR*, we have also identified *MAPKAPK3*, *ARNT*, and *ANGPTL1* as potential therapeutic targets in wet AMD. Our findings revealed that genetically predicted higher expression levels of *PLA2G7, MAPKAPK3, ARNT* were linked to an increased risk of AMD and *ANGPTL1* was associated with a decreased risk of AMD. We found that Trebananib can treat some tumors by targeting Angiopoietin-1 (ANGPT1) and Angiopoietin-2 (ANGPT2) (Leary et al., 2017). Drugs acting on the *ARNT* target, such as HIF-1alpha, can inhibit neovascularization in tumors by affecting the fibroblast growth factor receptor (FGFR), Hypoxia-inducible factor 1 alpha (HIF-1A), and VEGF (NCT00880672).

## Discussion

Our research utilized extensive genomic datasets and a novel multi-omic approach to examine immune and metabolic factors linked to AMD subtypes. We have identified immune cells, metabolites, and inflammatory proteins causally linked to AMD, and provide evidence of a protective mechanism against wet AMD involving the androgen-IL10RA-CD16+ monocyte axis, confirmed through mediation MR analysis and scRNA-seq. Additionally, we found the inflammatory response, TNFα and Notch signaling pathways to be critical in progressing early AMD to CNV, while BA metabolism is significant in dry AMD. Through ML and genomic MR, we have identified genes associated with AMD subtypes, noting a notable causal relationship in HLA-associated genes for both dry and wet AMD. We also highlight potential genetic drug targets, including *MTOR, PLA2G7, MAPKAPK3, ANGPTL1,* and *ARNT*, as candidate drugs for AMD treatment.

In our MR analysis, we have found significant causal relationships between the PE pathway and all subtypes of AMD, consistent with previous research findings (Gugiu et al., 2006). The development of AMD is significantly influenced by the PE pathway, as seen in the accumulation of bisretinoid compounds like A2-GPE and all-trans-retinal dimer-PE (Sparrow et al., 2012). These compounds, related to PE metabolism, accumulate in RPE cells, contributing to disease progression (Ueda et al., 2016). Epidemiological studies indicate that women appear to have a higher incidence of AMD than men (Rudnicka et al., 2012; Smith et al., 1997). A study has shown that serum androgen levels are inversely related to the severity of AMD, with AMD patients having significantly lower dehydroepiandrosterone sulphate levels compared to the normal group (Tamer et al., 2007). In another study, treatment with the male sex hormone antagonist 5α-reductase inhibitor was linked to a higher risk of macular abnormalities, hinting at the protective role of androgens in the macular region (Shin et al., 2020). In keeping with these observations, we have identified a significant causal relationship between the androgen pathway and wet AMD. IL-10, which is an anti-inflammatory cytokine, controls inflammation by binding to IL-10 receptor, activating anti-inflammatory genes via the JAK1/STAT3 pathway in hematopoietic cells such as lymphocytes and macrophages (de Souza et al., 2024). Our proteomic MR analysis discovered a significant negative causal relationship between IL10RA and wet AMD, while our mediator MR analysis revealed that three androgen-related metabolites are likely to inhibit wet AMD by enhancing IL10RA expression, suggesting the androgen pathway could play a protective role in wet AMD through the regulation of IL-10 pathway. Through scRNA-seq, we have found that pDCs and CD16+ monocytes exhibited a strong androgen response, particularly in wet AMD, with CD16+ monocytes showing elevated IL10RA expression. CD16+ monocytes are known to be involved in the pathogenesis of autoimmune and chronic inflammatory diseases (Hector and Sørensen, 2017; Poitou et al., 2011). Interestingly, we also found a significant causal relationship with AMD in our immunome MR analysis. Another clinical study found a significant association between CD16+ monocytes and refractory wet AMD (Lin et al., 2024). Coupled with scRNA-seq results, we hypothesize that the androgen - IL10RA - CD16+monocytes pathway is likely to be a critical immunometabolic pathway in wet AMD. We provide convincing evidence that immune cells, metabolites and inflammatory proteins are playing a significant role in pathogenesis, offering new angels for targeted therapeutic approaches in wet AMD.

After confirming the significant causal relationship of the androgen pathway in wet AMD and the immunometabolic mechanisms, we comprehensively mapped transcriptomic landscapes across 50 key pathways for all groups under study. First, we uncovered notable upregulation of angiogenesis in mixed and CNV groups, matching clinical observations and previous studies (Bressler, 2009). Furthermore, we have highlighted unique pathways like the complement, fatty acid and interferon pathways that are distinctly associated with specific AMD subtypes such as GA, when compared to controls (Armento et al., 2021b; Haines et al., 2005; Ren et al., 2022). These findings, which are consistent with previous studies, further validate our findings. Second, the inflammatory response, in particular, TNFα and Notching signaling pathways showed increased activity in early AMD and CNV. The activation of the Notching pathway has been found to decrease CNV lesions by influencing retinal vascular growth, and its inhibition exacerbated CNV, identifying crucial Notch targets in vascular homeostasis and CNV development (Ahmad et al., 2011). In another study, inhibiting the Notch pathway proved to be an effective method for suppressing retinal fibrosis in late-stage wet AMD (Fan et al., 2020). TNF-α is thought to directly contribute to CNV enhancement and has been shown to exert its effect by augmenting the inflammatory response through other signaling pathways (Papadopoulos, 2024). It should be emphasized that previous studies mainly focused on the Notch and TNF-α pathways in later stages of AMD, with limited data on early AMD. Our findings indicate that both pathways are highly expressed in early AMD, akin to the CNV group, pointing towards their crucial role in advancing early AMD to CNV. Thus, targeting these pathways could potentially prevent CNV development. Furthermore, Heme metabolism imbalance increases oxidative stress, a key factor in dry AMD development, damaging retinal cells and leading to disease progressing (Synowiec et al., 2012; Welch et al., 2002). Our study found elevated Heme expression in dry AMD, but not wet AMD, pointing towards oxidative stress as a significant pathogenic mechanism in dry AMD. Additionally, we found that the BA metabolism pathway is likely to play a significant role in dry AMD. Previous studies have indicated that BA is involved in the regulation of various retinal diseases, including retinitis pigmentosa and diabetic retinopathy (Win et al., 2021). BA plays a critical role in cholesterol catabolism and its involvement in AMD is not surprising, given that large cholesterol-rich deposits are a pathological hallmark (Omarova et al., 2012). Our research findings indicate higher expression of BA in dry AMD, suggesting their significant role in disease pathogenesis. As BA therapy has demonstrated positive effects in animal models of CNV, it could potentially serve as a therapeutic target for dry AMD (Warden et al., 2020). Furthermore, myogenin has been identified as a common mediator of FAP, TNC and GRP, and it could, therefore, be involved in AMD by regulating these target genes (Zhao et al., 2017). We have also uncovered significant expression of myogenesis in AMD, in keeping with previous work showing its association with various immune cell functions, aging, and inflammation (Tidball et al., 2021). However, its specific role in AMD requires further investigation.

To better understand the genes related to immunometabolic mechanisms in AMD, we used ML for gene analysis of scRNA-seq. Additionally, we employed genome-wide MR to identify AMD-related genetic loci and confirm their regulatory roles on associated immunometabolic pathways. First, the causal association of *CXCL9, CX3CR1 and ZNF683* with wet AMD were validated by both ML and genome-wide MR methods, strengthening their causal association with AMD. Second, we observed a significant depletion of certain immune cells and a reduction in cell-cell communication, particularly affecting monocytes, DCs and T cells in CNV patients compared to those with GA. These disruptions in peripheral immune interactions, including variations in key signaling pathways and ligand-receptor pairs, indicate a more pronounced pro-inflammatory state in CNV, suggesting underlying immunological differences between the AMD subtypes. Third, HLA antigens are normally expressed in the human eye, including in retinal microglia and uveal structures, but in AMD, there is intense HLA-DR and increased HLA class II immunoreactivity associated with drusen formation and CNV (Bakker et al., 1986; Penfold et al., 1997). The HLA system, which is crucial for innate and adaptive immune responses, has been the subject of intense interest in relation to the pathophysiology of AMD with several studies reporting a correlation between HLA genetic variants and AMD. However, there are also contradictory viewpoints with other studies reporting no association between HLA class II genes and AMD (Goverdhan et al., 2005; Goverdhan et al., 2008; Hageman et al., 2001; Mullins et al., 2000; Pappas et al., 2015; Villegas Becerril et al., 2009). In our study, we observed marked dysregulation in the interactions between HLA-DR-CD4 in myeloid DCs or monocytes and CD4+ T cells, emphasizing the strong influence of HLA-DR in the development of AMD. Moreover, our comprehensive genomic MR analysis provides compelling evidence of a significant causal link between *HLA-DQB1, HLA-DQB1-AS1, HLA-DQB2,* and *HLA-DRB6* and both dry and wet forms of AMD. *HLA-DQA1, HLA-DQA2,* and *HLA-DRB5* also showed significant causal relationships with dry AMD, highlighting the link between HLA class II genes and AMD. Next, we selected a series of immune cells that highly express HLA-DR and have a causal relationship with AMD to conduct a mediation MR analysis. This analysis revealed that monocytes and DCs with high HLA-DR expression act as mediators, linking various metabolites to AMD and underscoring the critical function of HLA-DR in its immunometabolic regulation.

Based on MR analysis, the *MTOR* gene is associated with both dry and wet AMD. The mammalian target of rapamycin (MTOR) is involved in cell growth, survival, autophagy and metabolism, operating through two complexes: MTORC1 and MTORC2 (Laplante and Sabatini, 2009). Dysregulation of the MTOR signaling pathway plays a role in a variety of diseases, including cancer, neurodegenerative disorders and diabetes mellitus (Griffin et al., 2005; Masui et al., 2013; Mossmann et al., 2018). The neuroprotective effects of MTOR inhibitors could offer a new treatment approach for AMD, targeting shared pathways like oxidative stress that lead to retinal neuronal death (Armstrong et al., 2022). *PLA2G7* is a key regulatory gene for understanding how adipose tissue metabolism influences immunometabolic responses (Candels et al., 2022). Rilapladib has demonstrated anti-inflammatory, antioxidant, and vascular-enhancing effects by targeting Phospholipase A2 (PLA2) and Platelet-activating factor acetylhydrolase (PLA2G7) in the treatment of Alzheimer’s disease (AD) (Maher-Edwards et al., 2015). Darapladib, another PlA2 inhibitor, has also been found to be beneficial in the treatment of diabetic macular edema (Staurenghi et al., 2015). Currently, the second-generation PLA2 inhibitor, SNP318, is undergoing a clinical trial for the treatment of age-related neurodegenerative diseases (NCT05792163). PLA2 and VEGF interact to regulate the proliferation and migration of RPE cells, which likely contribute to the pathophysiology of AMD (Kehler et al., 2012). Therefore, we speculate that PLA2 inhibitors are promising candidate drugs for AMD that could be explored in future clinical trials. *MAPKAPK3*, situated on chromosome 3 and encoding a 382 amino acid serine/threonine kinase, is activated by p38 kinase in response to stress. It shares 75% sequence identity with its close relative, *MAPKAPK2*, highlighting their key roles in response to stress factors such as DNA damage, oxidative and osmotic stresses (Mayer et al., 2001; Ronkina et al., 2011). *MAPKAPK3* is highly expressed in RPE cells, in keeping with its close association with the normal physiological functions of the retina (Meunier et al., 2016). In hereditary retinal diseases, such as patterns macular dystrophy, mutations in *MAPKAPK3* lead to abnormalities in RPE and Bruch’s membrane, contributing to the development of CNV, macular atrophy and retinal fibrosis, which show strong similarities to late-stage wet AMD. Our MR analysis found a causal relationship between the *MAPKAPK3* and wet AMD, suggesting that *MAPKAPK3* could be a potential drug target. Human angiopoietin-like proteins (ANGPTLs), a family of 8, play key roles in regulating metabolism, stem cell expansion, inflammation, angiogenesis and wound healing, reflecting their broad biological significance (Sodhi et al., 2019). Dysregulations in these responses contribute to the pathogenesis of AMD and other eye diseases (Yang et al., 2018). ANGPTL2 and ANGPTL4 have been shown to play important roles in the development of AMD (Hirasawa et al., 2016; Sodhi et al., 2019), but *ANGPTL1* can inhibit angiogenesis by targeting the VEGFA/VEGFR2/Akt/eNOS pathway (Wang et al., 2024). Our MR analysis indicated that *ANGPTL1* has a protective effect, suggesting that a potential therapeutic role in AMD. As a vital part of the hypoxia-inducible transcription factors (HIF) complex, aryl hydrocarbon receptor nuclear translocator (ARNT) significantly influences wet AMD by modulating angiogenic factors like VEGF and their receptors (Okabe et al., 2014). This regulation promotes pathological angiogenesis and vascular permeability, driving the progression of wet AMD (Campochiaro, 2013). In our scRNA-seq metabolites enrichment, we also found that hypoxia-related pathways are significantly upregulated in wet AMD. This underscores ARNT’s pivotal role in the hypoxia-responsive mechanisms and potential therapeutic target in wet AMD (Cammalleri et al., 2016).

Our study has notable strengths. We first analyzed large GWAS datasets of early, dry, wet and mixed AMD subtypes using MR, and categorized 43 AMD samples similarly for scRNA-seq analysis. This allowed us to profile the immunometabolic characteristics ofAMD subtypes, perform genetic analyses, and identify drug targets using both MR and scRNA-seq. We also used extensive datasets to examine the impact of immune cells/traits, circulating metabolites, and inflammatory proteins on AMD subtypes through a comprehensive multiomics MR approach. Specifically, we linked specific metabolites identified as causally related to AMD with their metabolic pathways, immune cells, and inflammatory factors using a novel combinatorial approach with mediated MR and scRNA-seq. We have highlighted the role of the androgen pathway - IL-RA10 - CD16+ monocyte axis in wet AMD, which to our knowledge, has not been implicated previously. By combining ML and genome-wide MR analyses, we have identified genes linked to the pathogenesis of AMD, in particular, a causal link with several HLA-associated genes. We have also generated a list of potential drug targets that could act on these pathways. Our study provides a comprehensive analysis of the immunometabolic mechanisms of AMD subtypes using a multiomics MR and scRNA-seq approach, offering valuable insights for understanding AMD pathogenesis and identifying treatment targets.

Our study has several limitations. First, effect sizes from MR analyses should be interpreted with caution. This is because the MR estimate is better interpreted as a test statistic for a causal hypothesis rather than the expected impact of a clinical intervention at a specific point in time (Burgess et al., 2019; VanderWeele et al., 2014). Second, although we did not observe evidence of pleiotropy for the causal association by different MR approaches, there is still a possibility that variants used in the MR confer a risk of AMD through a pleiotropic pathway (Davies et al., 2018; Slob and Burgess, 2020). Third, our genome-wide MR analyses only used cis-eQTLs to predict gene expression, while trans-eQTLs and other elements also regulate gene expression. Furthermore, our study focused on participants of European ancestry restricting the wider applicability of its conclusions to those of non-European heritage. The critical value of multiancestry MR comparisons is made clear by the current shortage of data concerning non-European ancestries. This underscores the significance of conducting comprehensive and well-supported genetic research within diverse ancestries to enhance the generalizability of a particular set of findings (Zhao et al., 2022; Zheng et al., 2022). In addition, the eQTLs used in our genome-wide MR analysis are from the Whole Blood Cell Database, and AMD is an ocular disease so many genes associated with AMD may not be expressed in blood cell types, limiting the statistical power and precision of our genome-wide MR results. Lastly, our MR analysis used the GWAS dataset of AMD in a large population, whereas the scRNA-seq analysis used patients with AMD collected in a clinical trial, and there may be differences in enrollment criteria and disease severity between the MR group and the single-cell RNA sequencing group, even in the same subtype of AMD.

In conclusion, our study identified immunometabolic signatures, related genes and drug targets for particular AMD subtypes. The androgen - IL10RA - CD16+ monocytes axis in particular seems to be important to the development of wet AMD. We identified similarities and differences between dry and wet AMD and explored some pathways that could drive the progression from early to advanced AMD. We validated with both ML and genome-wide MR methods a causal association between *CXCL9*, *CX3CR1* and *ZNF683* with wet AMD. Importantly, five genes, including *MTOR, PLA2G7, MAPKAPK3*, *ANGPTL1* and *ARNT* are promising drug targets for AMD, which will need to be explored further to evaluate their effectiveness in AMD prevention and treatment. We propose the integrated analytical model detailed in our study as a generalizable tool that can be applied to dissect the pathophysiology of other complex diseases.

## Method

A study overview is presented in Figure. 1, including data sources, study designs, methods, and follow-up analyses.

### Immune Cells/traits, Inflammatory Proteins and Metabolites to AMD

For the exposures GWAS, we used the GWASs for a total of 731 cell traits assessed in a general population cohort of 3,757 Sardinians, 1400 metabolites assessed in a cohort of 8,299 unrelated European ancestry individuals and 91 inflammatory proteins measured using the Olink Target platform in 14,824 participants (Table S1). For the outcome, we used the GWASs from Finland cohort in dry AMD, wet AMD, Mixed AMD and Early AMD from European cohort (Table S1).

For two-sample MR, the effect of exposures on AMD subtypes was assessed using the IVW method with a random-effects model in TwoSampleMR v.0.5.6 (Hemani et al., 2018) (https://mrcieu.github.io/TwoSampleMR/). The instrumental variables for the exposures were defined as genome-wide significant and independent SNPs (*P* < 5 × 10^−8^; r^2^ < 0.001, with a clumping window of 10 Mb). Data harmonization and MR analyses were executed via TwoSampleMR version 0.5.6. Findings that yielded a Q-value below 0.05 were classified as having substantial heterogeneity, indicating significant variation among the results (Tian et al., 2024) (Table S14∼17). To assess the presence of directional pleiotropy, the MR-Egger intercept test was conducted through the ‘mr_pleiotropy_test()’ function. Evidence of directional pleiotropy was identified if the MR-Egger intercept was statistically different from zero, indicated by a P value of less than 0.05. It’s important to highlight that the existence of moderate heterogeneity does not contravene the MR principle that presupposes the absence of directional pleiotropy, provided that the horizontal pleiotropic effects are balanced (Cho et al., 2020) (Table S18∼21). In the reverse MR analysis, which evaluated the impact of AMD on immune cells/traits, inflammatory proteins, and metabolites, SNPs associated with AMD subtypes were utilized as the instrumental variables for the exposures. The IVW method was employed, and in cases where only a single SNP was available, the Wald ratio method was applied. Findings yielding a *P* value less than 0.05 were deemed statistically significant (Table S18∼21). To assess statistical power, *F-* statistics were calculated as previously described using the following formula: 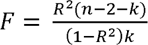 where: R^2^ = proportion of variance in the exposure trait and k = number of instrumental variables (Pierce et al., 2011) (Table S2∼13). We used a relaxed FDR (Adjusted *P*-value by the Benjaminiand Hochberg method) threshold of 0.2 because of the relatively lower SNPs sizes of most of single immune cells, metabolites and inflammatory proteins (Mavromatis et al., 2023)

### Enrichment of Metabolites in AMD

We performed enrichment analysis on metabolites with significant effects (*P*-values < 0.05) across AMD subtypes, mapped against a curated reference of metabolic pathways (Table S39). Enrichment ratios were calculated by dividing the count of group-specific metabolites within each pathway by the total count of metabolites cataloged in that pathway from the reference set. We employed the hypergeometric test to determine the statistical significance of enrichment for each pathway, which considers the size of both the reference set and the group-specific set. In recognition of the multiple hypothesis testing issue inherent to such analyses, we adjusted the resultant p-values. The Benjamini-Hochberg correction procedure was utilized to control the false discovery rate, ensuring a robust statistical framework for identifying significantly enriched pathways.

### Mediation Analysis

Through mediation MR, we explored the causal relationships between immune cells/traits, metabolites, and inflammatory proteins with AMD to identify potential immunometabolic pathways. Additionally, we were able to calculate the proportion of the mediating effect in the total effect (Table 26∼31).

First, we used androgen-related metabolites as the exposure, with immune cells and inflammatory proteins as the outcomes, to determine the causal relationship between the metabolites and immune cells/inflammatory proteins. The effect size from this step is denoted as β1. Second, using the immune cells/inflammatory factors that have a causal relationship with the metabolites identified in the first step as the exposure, and AMD subtypes as the outcome, we conducted a MR analysis, resulting in an effect size of β2. Then, multiplying β1 by β2 yields β12, which represents the effect size of this mediating pathway. The total effect size between androgen metabolites and AMD obtained from the two-sample MR analysis serves as the total effect size. Dividing β12 by the total effect size gives the proportion of this mediating effect. We also analyzed the pleiotropy, heterogeneity, and conducted reverse Mendelian Randomization for the mediating effects to ensure the accuracy of our results. To explore more potential immunometabolic mechanisms, in addition to androgen metabolites, we conducted MR analyses on the interrelationships among immune cells, metabolites, and inflammatory proteins. This allowed us to identify through which interactions they can affect AMD and exert regulatory effects on AMD.

### Identify causal genes associated with AMD

First, we employed MR analyses to identify potential causal genes of AMD. We use the eQTLs of the whole blood from “GTEx_Analysis_v8_eQTL_EUR” which are Cis-eQTLs mapped in 838 postmortem donors from European-American subjects. We employed cis-eQTLs within 100Kb of gene probes as genetic instruments for expression levels in whole blood, focusing on their impact on dry and wet AMD. We employed cis-eQTLs within 100Kb of gene probes as genetic instruments for expression levels in whole blood, focusing on their impact on dry and wet AMD. These instruments were selected based on stringent criteria: F-statistics > 10 for instrument strength, association with gene expression (p < 1.0 x 10^-5), and Steiger filtering to exclude reverse causality (Hemani et al., 2017). We conducted MR analyses to assess the potential causal effects of gene expression levels on dry and wet AMD respectively. After MR analysis, we identified 31 potential causal genes with robust signals (*P* < 0.000015, FDR test by *P* < 0.05/3230) in dry AMD and 29 potential causal genes in wet AMD (Table S32∼33). Subsequently, we analyzed genes that have a causal relationship with both dry AMD and wet AMD, discovering that there are 9 genes common to both conditions. We found that many HLA-related genes have a significant causal relationship with both dry AMD and wet AMD. Therefore, through mediator MR analysis, we explored how a series of immune cells related to HLA exert a regulatory effect on AMD through their interactions with metabolites or inflammatory proteins.

### Identify Drug Targets for AMD Subtypes

Based on pathogenic genes identified in previous MR analyses, we explored potential drug targets by querying Open Targets and the Therapeutic Target Database. We investigated diseases with clinical traits, pathogenesis mechanisms similar or identical to AMD, such as macular malnutrition, neurodegenerative diseases, and neovascular-related diseases. We discovered that genes previously confirmed to be related to these diseases share some pathogenic genes with AMD. These genes, which can affect diseases similar or related to AMD, are hypothesized to be potential drug targets specifically influencing AMD. Consequently, we conducted an extensive drug search for all pathogenic genes identified in this study, with particular focus on these shared pathogenic genes. To gather information on potential target genes and their clinical annotations, we utilized two primary sources: the Therapeutic Target Database and Open Targets. TTD provides valuable information about therapeutic protein and nucleic acid targets, the specific diseases they are associated with, and the drugs related to these targets. Open Targets documents the relationships between genes and diseases, scoring each category of evidence to further confirm the feasibility of these pathogenic genes as drug targets (Table 2).

### ScRNA-seq Data Processing

The scRNA-seq datasets, encompassing blood samples from both healthy individuals and patients diagnosed with AMD, are made available to the public domain through the Gene Expression Omnibus (GEO) repository, through the accession code GSE222647 (Lin et al., 2024). The initial phase of quality control was executed by applying a set of criteria for cell selection: specifically, only those cells were considered that exhibited a gene expression count ranging from more than 400 to less than 8,000 and an UMI (Unique Molecular Identifier) count that fell between 600 and 120,000. Cells were also excluded if they presented an excessive mitochondrial gene expression, defined as exceeding 10% of their total gene count. Following this, the retained data matrix was subjected to normalization, utilizing the ‘LogNormalize’ technique as implemented in the Seurat R package (version 3.2.2) (Stuart et al., 2019), with a normalization factor set at 10,000. This process also included the regression of variables such as ‘Percent.mito’ and ‘nCount_RNA’ during the data scaling phase to remove confounding variables. Subsequent to these preparatory steps, the harmonization of all sample datasets was achieved through the application of the harmony R package (version 1.2.0), effectively minimizing batch effects and ensuring comparability across the study.

### Unsupervised Clustering and Annotation of Cell Types

For the purpose of identifying highly variable genes that facilitate the unsupervised clustering of cells, the Seurat package (version 5.0.2) was utilized. This was followed by performing a principal component analysis (PCA) focused on the top 2,000 variable genes to uncover underlying patterns. To select the principal components (PCs) crucial for further analyses, the ElbowPlot tool within Seurat was used, leading to the identification of 20 significant PCs. These selected components then served as the basis for advanced dimensionality reduction techniques, specifically through the generation of visual mappings using either Uniform Manifold Approximation and Projection (UMAP) or t-Distributed Stochastic Neighbor Embedding (t-SNE), facilitated by Seurat’s RunUMAP and RunTSNE functions, respectively. The approach for annotating cell types was in line with methodologies previously described (Lin et al., 2024).

### Gene Set Score Calculation

To quantify the activity of specific signaling pathways in the context of AMD, curated gene sets named “Hallmark” from the Molecular Signatures Database (MsigDB) were employed (http://www.gsea-msigdb.org/gsea/msigdb/index.jsp). The AddModuleScore function in Seurat (version 5.0.2) was utilized for the computation of pathway activity scores, applying the function’s default parameter settings. Averages of these module scores were then computed across different disease states, delineating the variance in pathway activity across the AMD progression spectrum. Statistical analysis was conducted to compare pathway activities between the control group and various stages of AMD, employing the Wilcoxon Rank Sum test, followed by adjustment for multiple comparisons through the Benjamini-Hochberg method.

### Gene Set Enrichment Analysis

For our study, the identification of differentially expressed genes (DEGs) was performed using the Seurat toolkit (version 3.2.2), leveraging the FindMarkers function to analyze variations across groups. Following the identification of DEGs, we proceeded with gene set enrichment analysis (GSEA) (Subramanian et al., 2005) to investigate the biological significance of these genes in relation to known biological pathways and processes. This analysis was carried out using the clusterProfiler package (version 3.18.1) (Yu et al., 2012). The reference gene set employed for the GSEA was the ImmuneSigDB gene sets from the Molecular Signatures Database (MsigDB), chosen specifically for its focus on immune-related pathways.

### Cell-cell Communication Analysis

Our analysis focused on cell-cell communication within AMD’s microenvironment, employing CellChat (version 1.1.3) (Dimitrov et al., 2022) on scRNA-seq data from PBMCs of healthy controls and AMD patients, specifically those with geographic atrophy (GA) and choroidal neovascularization (CNV). For each group, CellChat was used to construct detailed communication networks, starting with the identification of overexpressed genes and their projections onto known protein-protein interactions (PPIs). The computation of communication probabilities between cell types, both at the individual interaction and pathway levels, allowed for a quantitative assessment of signaling dynamics, which was further aggregated to elucidate dominant communication trends. Subsequent stages of the analysis involved the comparison of interaction strengths across different AMD phenotypes, illustrating differences in ligand-receptor interaction counts and weights.

### Visualization of Gene Expression and Cellular Proportions

The visualization of specific gene set scores, gene expression profiles, and cell proportions was accomplished using a suite of graphical representations including heatmaps, violin plots, and radar charts. These were generated by employing the pheatmap package (version 1.0.12), which can be accessed at (https://cran.r-project.org/web/packages/pheatmap), in addition to the dittoSeq package (version 1.2.4) (Bunis et al., 2020), and the ggplot2 package (version 3.5.0) (Wickham and Wickham, 2016). The data underwent an automatic scaling process facilitated by the default settings inherent to these packages, ensuring a standardized approach to data representation and enhancing the interpretability of the visualizations.

### Statistics

#### MR analysis

MR analysis leverages numerous SNPs as instruments alongside multiple gene expression traits as exposures, allowing for a multifaceted approach to the study. This analysis adheres to three fundamental MR principles: firstly, the relevance principle, ensuring that genetic predictors are strongly associated with the exposure in question; secondly, the exchangeability principle, which guarantees that the relationship between instruments, exposures, and outcomes remains unconfounded; and thirdly, the exclusion restriction principle, stating that the instruments influence the outcome solely via the exposure under investigation. Our reporting complies with the STROBE-MR guidelines, which aim to enhance the clarity and completeness of Mendelian Randomization studies.

#### LD check

In GWAS and MR analyses, clumping is used to select independent genetic variants (instruments) by removing variants that are in strong LD with each other. This is done to ensure that each SNP used as an instrument in MR analyses represents a unique signal, reducing the risk of bias due to correlated predictors. In our study, the instrumental variables for the exposures were defined as genome-wide significant and independent SNPs (P < 5 × 10^−8^; r2 < 0.001, with a clumping window of 10 Mb).

#### scRNA-seq

The determination of differentially expressed genes (DEGs), execution of GSEA, and the assessment of cell-cell communication within scRNA-seq datasets were performed by extracting p-values through the default statistical frameworks provided by the respective R analytical packages. For the comparison of gene set scores among different cell groups, the ggpubr package (version 0.6.0), which can be found at (https://CRAN.R-project.org/package=ggpubr), was utilized.

## Data availability

Immune cell trait GWAS summary statistics used for MR: https://gwas.mrcieu.ac.uk/, GCST 90001391 through GCST-90002121; Metabolites GWAS summary statistics used for MR: https://gwas.mrcieu.ac.uk/, GCST90199621 to GCST90201020; eQTLgen whole blood eQTL data.

used for MR of the genome: https://www.gtexportal.org/home/downloads/adult-gtex/qtl; Inflammatory proteins GWAS summary statistics used for MR: https://gwas.mrcieu.ac.uk/, GCST90274758 to GCST90274848; Dry AMD, Wet AMD, Early AMD and Mixed AMD GWAS summary statistics used for MR: https://gwas.mrcieu.ac.uk/, finn-b-DRY_AMD, finn-b-WET_AMD, ebi-a-GCST010723 and finn-b-H7_AMD. Androgens and IL10 receptor GWAS used for two sample MR: https://gwas.mrcieu.ac.uk/, met-a-747, met-a-460, met-a-478, met-a-627, prot-a-1465. The scRNA-seq datasets are made available to the GEO with the accession code GSE222647.

## Acknowledgement

The authors express their gratitude to all nonauthor study investigators, sites.

XC is supported by Cambridge International PhD Scholarship, QZ is supported by Cancer Research UK Cambridge Centre PhD Studentship.

P.Y.-W.-M. is supported by an Advanced Fellowship Award (NIHR301696) from the UK National Institute for Health Research (NIHR); he also receives funding from Fight for Sight (UK), the Isaac Newton Trust (UK), Moorfields Eye Charity (GR001376), the Addenbrooke’s Charitable Trust, Cambridge University Hospitals, the National Eye Research Centre (UK), the International Foundation for Optic Nerve Disease (IFOND), the NIHR as part of the Rare Diseases Translational Research Collaboration, the NIHR Cambridge Biomedical Research Centre (NIHR203312), and the NIHR Biomedical Research Centre based at Moorfields Eye Hospital NHS Foundation Trust and UCL Institute of Ophthalmology (NIHR203322). The views expressed are those of the author(s) and not necessarily those of the NHS, the NIHR or the Department of Health.

This work is supported by National Natural Science Foundation of China (82171085), National Natural Science Foundation of China (81900891).

## Authors Contribution

X.C. and Q.Z. conducted the statistical analysis of the data. Q.Z., X.C. and X.L., significantly contributed to the design of the study. X.C, Q.Z., D.W and P.Y.-W.-M. made significant contributions to the writing of this manuscript. All of the other authors critically reviewed the manuscript.

