## Supplementary figures and images for "Multiomic Screening Unravels the Immunometabolic Signatures and Drug Targets of Age-Related Macular Degeneration"

### Supplement Figures.pdf

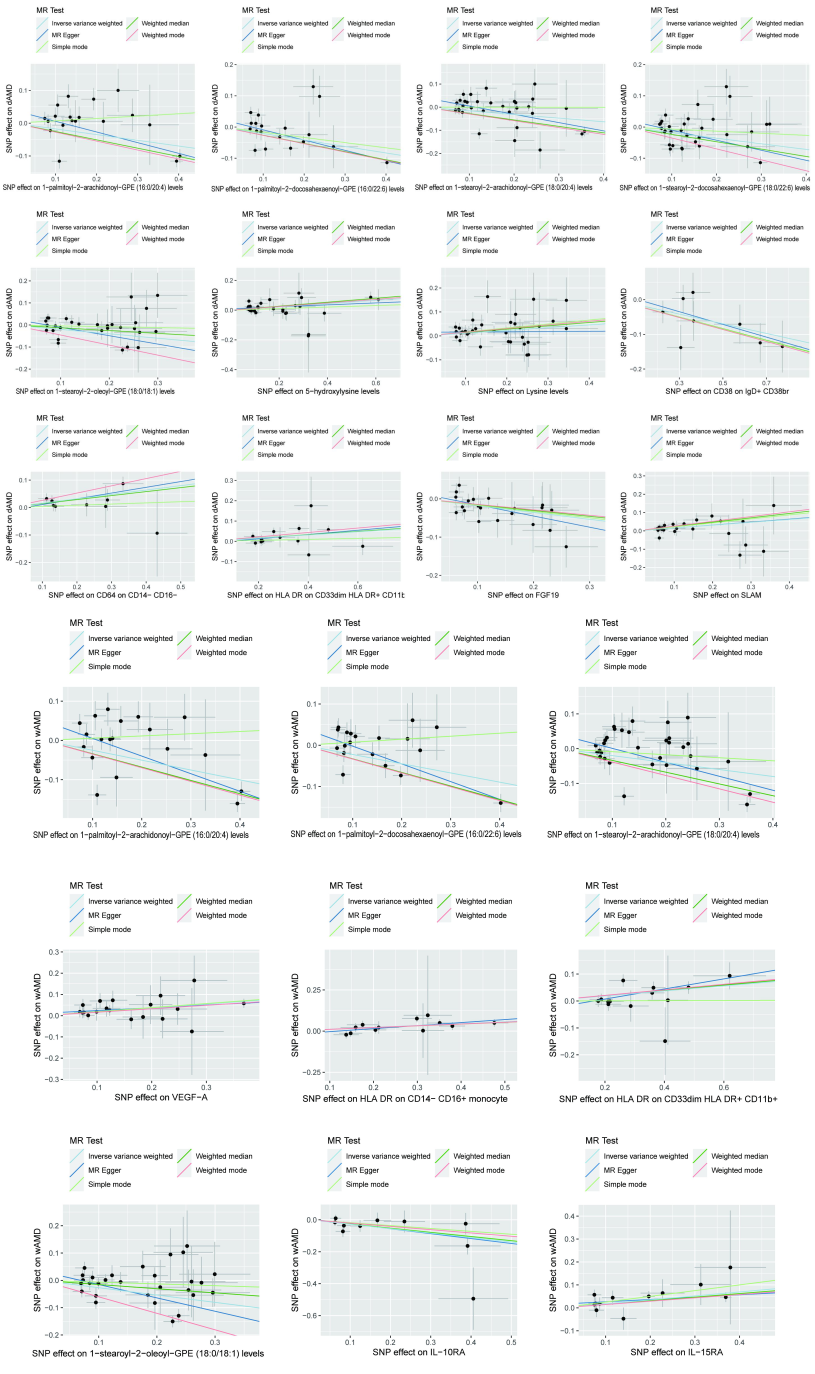

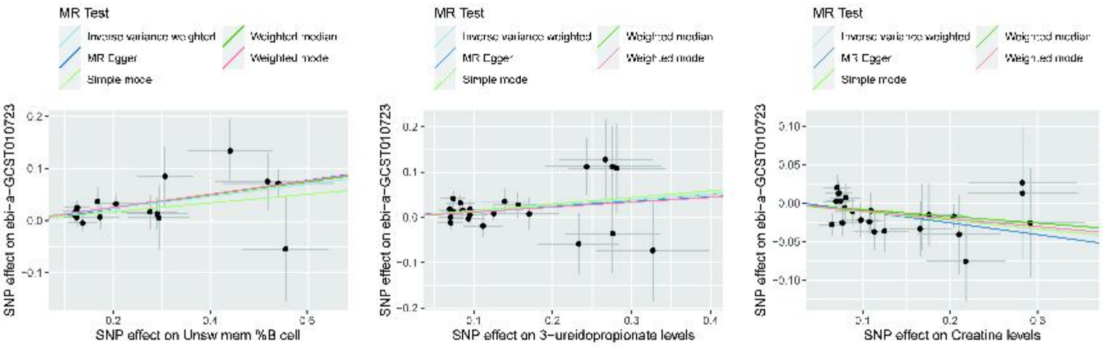

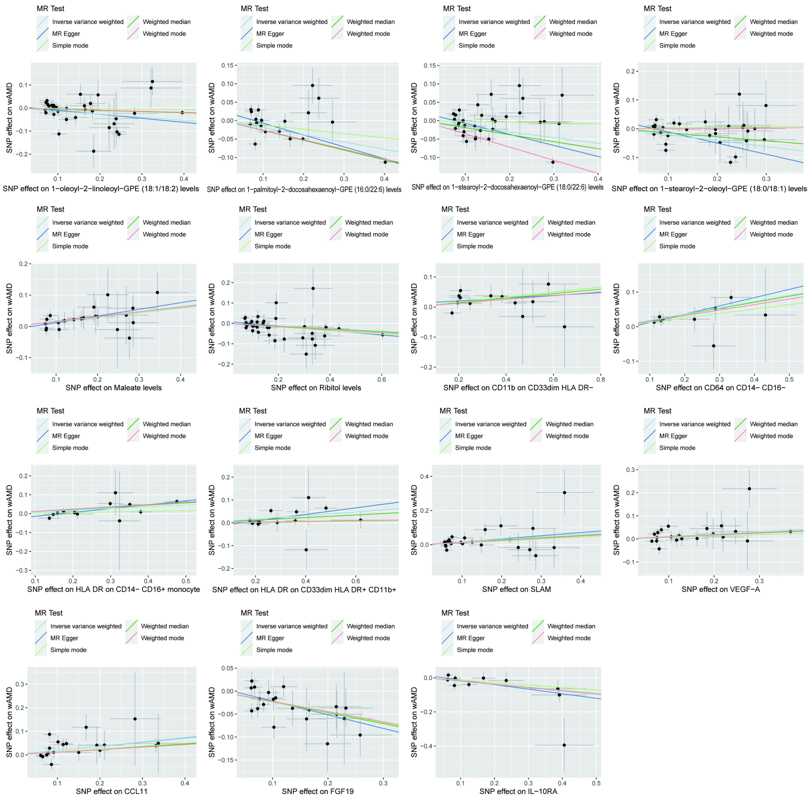

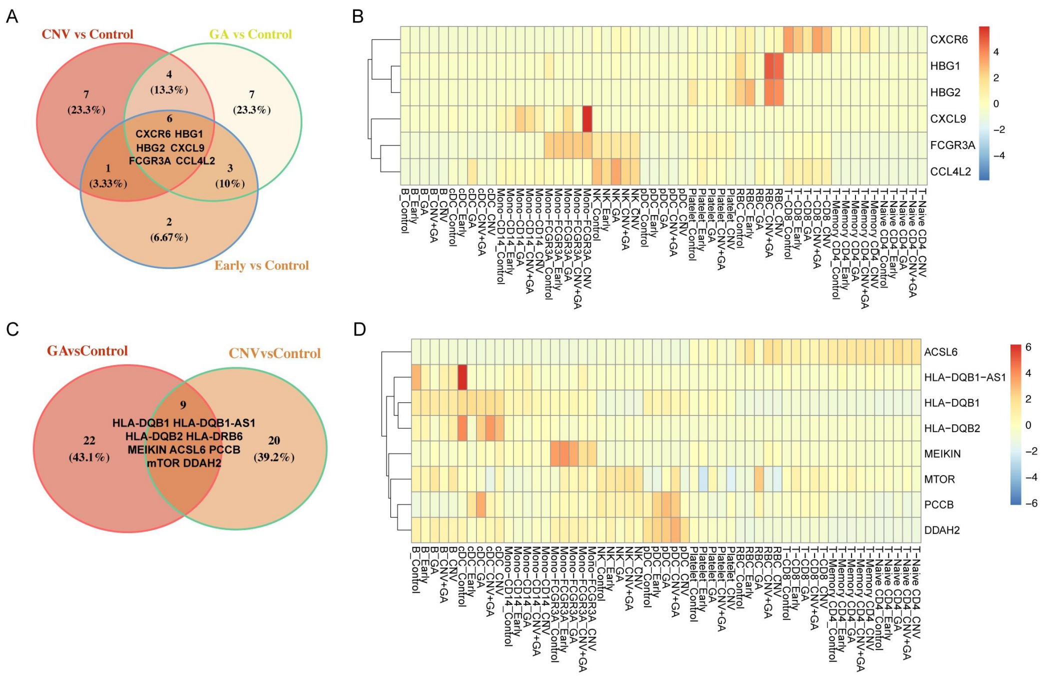

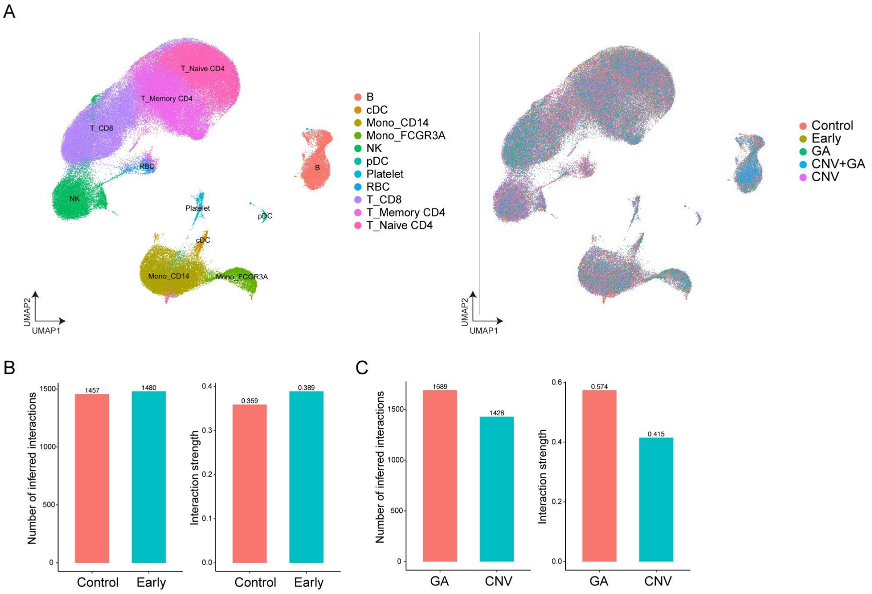
